# Low-frequency transcranial magnetic stimulation offers both immediate and delayed neuroprotection in neonatal hypoxia-ischemia model

**DOI:** 10.1101/2025.09.18.677186

**Authors:** Ivan Goussakov, Sylvia Synowiec, Alexander Drobyshevsky

**Affiliations:** Department of Pediatrics, Endeavor Health, Evanston, IL, 60201, USA

**Keywords:** perinatal brain injury, transcranial magnetic stimulation

## Abstract

Perinatal hypoxic-ischemic encephalopathy (HIE) is a leading cause of morbidity and mortality, and the current standard of care, therapeutic hypothermia, provides only partial neuroprotection. This study investigates the potential of low-frequency transcranial magnetic stimulation (LF-TMS) as a novel non-pharmacological adjunct therapy by targeting a key pathological mechanism of HIE: a persistent, pathological increase in glutamatergic synaptic transmission, or hypoxic long-term potentiation (hLTP).

Using a neonatal mouse model of hypoxia-ischemia, we administered a single session of LF-TMS shortly after the hypoxic event. We then evaluated its effects on synaptic function via slice electrophysiology and on brain injury volume using serial MRI. Our results show that hypoxia-ischemia induced a significant and lasting synaptic potentiation in the brain’s penumbral region. A single LF-TMS treatment successfully reduced this elevated glutamatergic response to control levels, suggesting a therapeutic mechanism similar to long-term depression (LTD) by regulating AMPA receptor redistribution.

Furthermore, LF-TMS provided significant neuroprotection, as demonstrated by a reduction in the volume of the ischemic core and penumbra 48 hours after the injury. LF-TMS did not alter excitability in sham-treated mice, confirming its safety as a targeted intervention for pathological conditions without affecting normal brain function. This study provides strong evidence that LF-TMS is a promising neuroprotective strategy that acutely and subacutely mitigates brain injury in a neonatal hypoxia-ischemia model.

## INTRODUCTION

Perinatal hypoxic–ischemic encephalopathy (HIE) is associated with high morbidity and mortality rates worldwide [1]. While therapeutic hypothermia remains the only approved treatment for near-term infants with neonatal hypoxia [2], it does not entirely protect an injured brain, especially in neonates with the most severe forms of hypoxic–ischemic injury [3]. his has created a critical need for adjunct interventions that can improve neuroprotection and extend the therapeutic window. Currently, several studies have failed to show a significant additional benefit in human clinical trials [3, 4].

The cellular pathogenesis of perinatal hypoxia involves primary and secondary energy failure, excitotoxicity, and oxidative stress. Our research and other published data suggest an additional mechanism: a pathological increase in glutamatergic transmission [5] due to hypoxic synaptic potentiation [6], which contributes to excitotoxicity and energy failure and can be targeted by therapeutic interventions. Previous pharmacological approaches targeting glutamatergic AMPA and NMDA receptors to mitigate excitotoxicity have been limited by significant challenges in achieving targeted and timely drug delivery to the precise sites of injury. Furthermore, the broad, non-specific nature of systemic drug administration poses a substantial long-term risk of interfering with critical neurodevelopmental signaling pathways, which can disrupt normal neuronal maturation.

Non-invasive brain stimulation (NIBS), such as low-frequency transcranial magnetic stimulation (LF-TMS), offers a promising adjunct non-pharmacological early intervention. It can reduce the pathologically increased glutamatergic synaptic transmission that is known to persist for days after perinatal brain injury [7]. This study investigates the neuroprotective effects of LF-TMS in a neonatal H-I model using postnatal day 9 mice, a developmental stage equivalent to human term newborns. We hypothesized that LF-TMS, when applied shortly after H-I, would mitigate the H-I-induced synaptic potentiation and reduce the volume of injured brain tissue.

## METHODS

The study has been approved by the Institutional Animal Care and Use Committees of Endeavor Health. The timeline for the experiments is shown in Figure 1.

**Figure 1.**
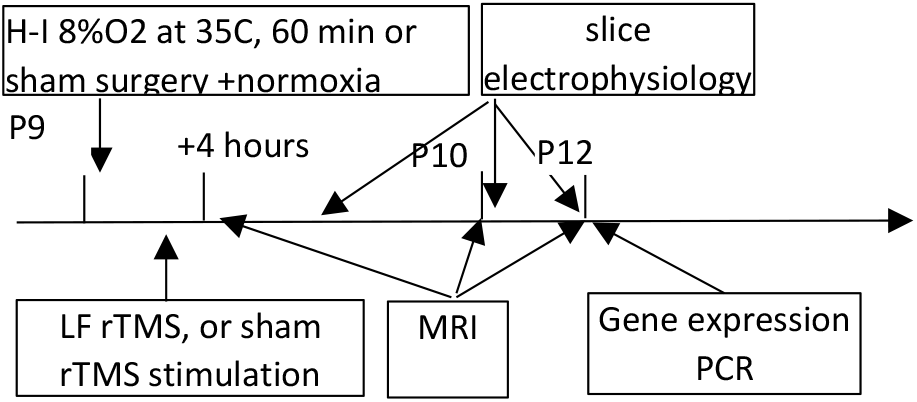
Timeline of the experiments.

### Murine model of near-term hypoxic-ischemic encephalopathy

The Vannucci model of neonatal hypoxic-ischemic (HI) brain injury was induced in postnatal day 9 (P9) C57Bl/6 mice [8], at a developmental stage commonly considered equivalent to the human term newborn. Briefly, pups were anesthetized with isoflurane, and the left common carotid artery was unilaterally isolated, cauterized, and cut. After a 2-hour recovery period with the dam, pups were exposed to systemic hypoxia in a humidified chamber maintained at 35.5°C within an incubator, with an inspired oxygen concentration of 8% balanced with nitrogen for 50 minutes. Sham-operated control animals underwent identical procedures of anesthesia and surgical incision but without carotid ligation and hypoxia. Sham animals were separated from their mothers and kept in the incubator for the same duration as the HI group, but without hypoxia.

### Transcranial magnetic stimulation

After a 30-minute recovery from hypoxia, pups were randomly divided into groups receiving TMS or sham treatment. TMS was administered to non-sedated mouse pups using the Neurosoft Transcranial Magnetic Stimulator (Soterix Medical Inc., Woodbridge, NJ) with a liquid-cooled 58x35 mm figure-of-eight TMS coil, optimized for rodent head stimulation [9]. The coil can deliver 2.6T at 80% power. TMS pulses were applied at 1Hz for 20 minutes, totaling 1200 pulses at 80% of the device’s maximum power. At this stimulation level, motor responses were visually observed in the control non-hypoxia group as hind limb and tail twitching (Supplementary video 1). TMS was delivered to groups of four pups placed in a custom holder with snug slots for soft immobilization, with their heads facing each other. The coil was positioned 5 mm above the center of the pups’ heads (Figure 1). Mice in the sham control groups (n=9, 7) were head-immobilized for the same duration but did not receive stimulation. During the session, pups rested on a temperature-controlled heated blanket connected to a water pump (Gaymar, USA).

### In-vivo MRI protocol

At 4, 24, 48, and 72 hours after H-I, serial MRI data were collected from all pups using a 9.4 T Bruker Biospec system (Bruker, Billerica, MA). Mice were sedated with isoflurane (Abbott, IL) inhalation, diluted in air to 3% for induction and 1.0% for maintenance. The animals’ respiration rate and skin temperature were monitored with a small animal physiological monitor (Model 1030, Small Animal Instruments, NY, USA). Body temperature was tracked using a rectal probe and maintained at 35 °C by blowing warm air. Animals were positioned prone in a cradle with their heads secured beneath the surface coil. Multi-echo T2-weighted images were acquired using an MSME sequence with TE/TR 10/2500 ms, 10 echoes, four signal averages, and an in-plane resolution of 0.12x0.12 mm. Twenty coronal slices with a thickness of 0.5 mm covered the entire brain. Slice pitch angles were set so that they were perpendicular to the plane of the localizer image, connecting the lowest point of the frontal and occipital cortex to facilitate slice co-registration of MR images and follow-up electrophysiology and tissue staining. [10]. T2 parametric maps were generated using Bruker Paravision 6.0 software by fitting an exponential decay function to signal versus echo times.

Automatic segmentation of injured brain tissue into two classes, ischemic core and penumbra, was performed using an automatic Hierarchical Region split procedure [3], implemented in Matlab. First, a mask of hyperintense brain areas indicating injured regions (edema) was manually outlined on each slice with ITK-SNAP 3.2 (itksnap.org). This step helped us avoid inaccuracies in the subsuquent segmentation caused by incorrect skull stripping and the selection of areas with a high contribution of cerebrospinal fluid, such as ventricles and the subarachnoid space. Then, the masked parametric T2 image was recursively split into two sub-images using the valley between these two modes in the T2 time histogram, and finally assigned to either the ischemic core class if the corresponding mode of the histogram was more than 56 ms or to the penumbra class otherwise. Voxel values with T2 times less than 33 ms were assigned to the uninjured brain tissue class. The thresholds for T2 values were set based on the pilot dataset from edematous brain regions at 24 hours after H-I that later developed into cystic lesions. Additionally, the volume of the brain tissue lost after H-I, excluding ischemic core and penumbra, was determined by subtracting the volume of the intact hemisphere from the injured hemisphere.

### Brain slice electrophysiology

A subset of mice underwent MRI, were deeply anesthetized with 3% isoflurane, and then decapitated. Their brains were quickly removed into ice-cold artificial cerebrospinal fluid (aCSF). The olfactory bulbs and cerebellums were removed, and the hemispheres ipsilateral to carotid ligation were marked with a notch. The brains were oriented for sectioning to match the MRI slice pitch angle as described above. Axial slices were cut with a thickness of 350 µm using a Leica vibrotome (VT1000S, Leica Biosystems). The slice preparation ACSF contained the following reagents in mM: 68 NMDG, 2.5 KCl, 1.25 NaH_2_PO_4_, 30 NaHCO_3_, 20 HEPES, 2 Thiourea, 25 D-glucose, 3 Na-ascorbate, 0.3 mL/L ethyl pyruvate, 0.5 CaCl_2_-2H_2_O, 10 MgSO_4_-7H_2_O, and 2 Kynurenic acid. The pH was adjusted to 7.3 with 1 N HCl after adding NMDG, then corrected to the final value after all other components were added. Final osmolality was 300±5 mOsm/kg H2O. NMDG was used for slicing as a substitute for NaCl to improve cell viability during slice preparation. Slices were transferred to a custom-made incubation chamber with a storage solution containing (in mM): 120 NaCl, 2.5 KCl, 1.25 NaH_2_PO4, 25 NaHCO_3_, 10 D-glucose, 2 Thiourea, 3 Na-ascorbate, 0.3 ml/L Na-Pyruvate, 2 CaCl_2_-2H2O, 2 MgSO_4_-7H_2_O, 2 Kynurenic acid, and 10 HEPES/NaOH, pH 7.4, with osmolality of 300±5 mOsm. The slices were kept on a nylon mesh in an incubation chamber and bubbled with carbogen for at least 1 hour before recording.

Slices were transferred to a recording chamber (model RC-24, Warner Instruments, Hamden, CT, USA) and continuously perfused with room-temperature, oxygenated aCSF bubbled with carbogen (pH 7.4) at a rate of 3 ml/min. The location of the recording sites was determined based on the locations of T2-hyperintense lesions on the corresponding MRI of each animal: 1) inside, 2) on the periphery, 3) remote (3-4 mm away from the hyperintense lesion), and 4) at the symmetrical sites of the contralateral hemisphere. Field excitatory postsynaptic potential (fEPSP) was recorded in cortical layer 3. Unipolar stimulus pulses with 0.1-ms width were delivered from a stimulus isolator, Iso-Flex (I.M.P.I., Israel). Stimuli to evoke fEPSP were applied once per minute during all recordings. A bipolar stimulation electrode, composed of 20 μm twisted platinum-iridium wires, was placed precisely at the border between the molecular layer and layer III, as visible under a 10x objective. The recording pipette, filled with extracellular solution, was positioned 250 μm from the cathode wire in the middle of layer III to minimize variability in fEPSP placement. Input-output curves were generated by increasing stimulus intensity in 50 μA increments from 0 to 650 μA and measuring fEPSP amplitudes. Paired pulse facilitation (PPF) was assessed using a pair of 50 μA stimuli separated by 75 ms intervals. PPF was calculated as the percentage increase in the amplitude of the second response relative to that of the first response of the pair. Electrical signals were amplified using a Multiclamp 700 B amplifier and digitized with a Digidata 1440 (Molecular Devices, Sunnyvale, CA, USA). The signals were recorded in current clamp mode using pCLAMP 10 software (Molecular Devices). All drugs were purchased from Sigma-Aldrich Inc. (MO, USA).

### RNA Isolation and Real-time PCR

After completing the MRI, mouse brains were harvested, and the cortices of the left and right hemispheres were separated. These were then individually frozen in liquid nitrogen and stored at -80°C before use. The tissue was homogenized on ice with Qiazol Lysis Reagent (cat. # 79306 Qiagen, Germantown, MD) for total RNA extraction. RNA quantity and quality were assessed using a NanoDrop spectrophotometer (ThermoFisher Scientific, USA). Next, 500 ng of isolated total RNA was used to synthesize cDNA with an RT2 First Strand Kit from QIAGEN. The cDNA was amplified via PCR using the QuantiTect Sybr Green PCR Kit (Qiagen, Germantown, MD) on an Applied Biosystems QuantStudio 7 Flex real-time quantitative PCR system. The total reaction volume was 25 μL, containing 1.0 μL of RT product (cDNA). Primers for Gria2 in the Sybr Green Assay were predesigned and ordered from Integrated DNA Technologies, Inc., USA. Primer sequences are listed in Supplementary Table 1. For each primer, the final concentration was 0.4 μM, with the annealing temperature adjusted accordingly. Gene expression was normalized to the housekeeping gene GAPDH and expressed as a fold change using the ΔΔCt method.

### Statistical analysis

To evaluate the main and interaction effects of recording location, time, and TMS, we conducted three-factor ANOVAs. Post-hoc Dunnett’s multiple comparisons test or Tukey’s HSD test was then used to identify specific group differences by comparing them to the values from the contralateral hemisphere. The evolution of volumes of segmented tissue classes on T2-weighted images was analyzed with a repeated measures ANOVA for each tissue class, followed by multiple comparisons between groups at each time point. Statistical analyses were performed using GraphPad Prism 10 (GraphPad Software, Inc., La Jolla, CA). A significance level of p ≤ 0.05 was applied.

## RESULTS

### fEPSP recordings distal and proximal to edema sites

The sites for extracellular field potential recordings on perfused brain slices were guided by individual animal MRI, which showed a strong hyperintense signal on T2-weighted scans, indicating cytotoxic edema in the ipsilateral hemisphere as early as 4 hours after H-I. At the earlier time point, 4 to 8 hours after H-I, we could not detect the injury sites on the perfused slices due to the unaltered appearance of cells on traditional differential interference contrast (DIC) imaging. At 24 hours after H-I, injury sites became visible under DIC contrast as opaque spots (Figure 2A, black arrow), surrounded by normally appearing cells (white arrow). A glass pipette for recording field potential was placed inside the edema lesions, 200-300 µm near the lesion’s border, 2-3 mm away from the lesion (Figure 2B-D), and on the contralateral hemisphere (not shown).

**Figure 3.**
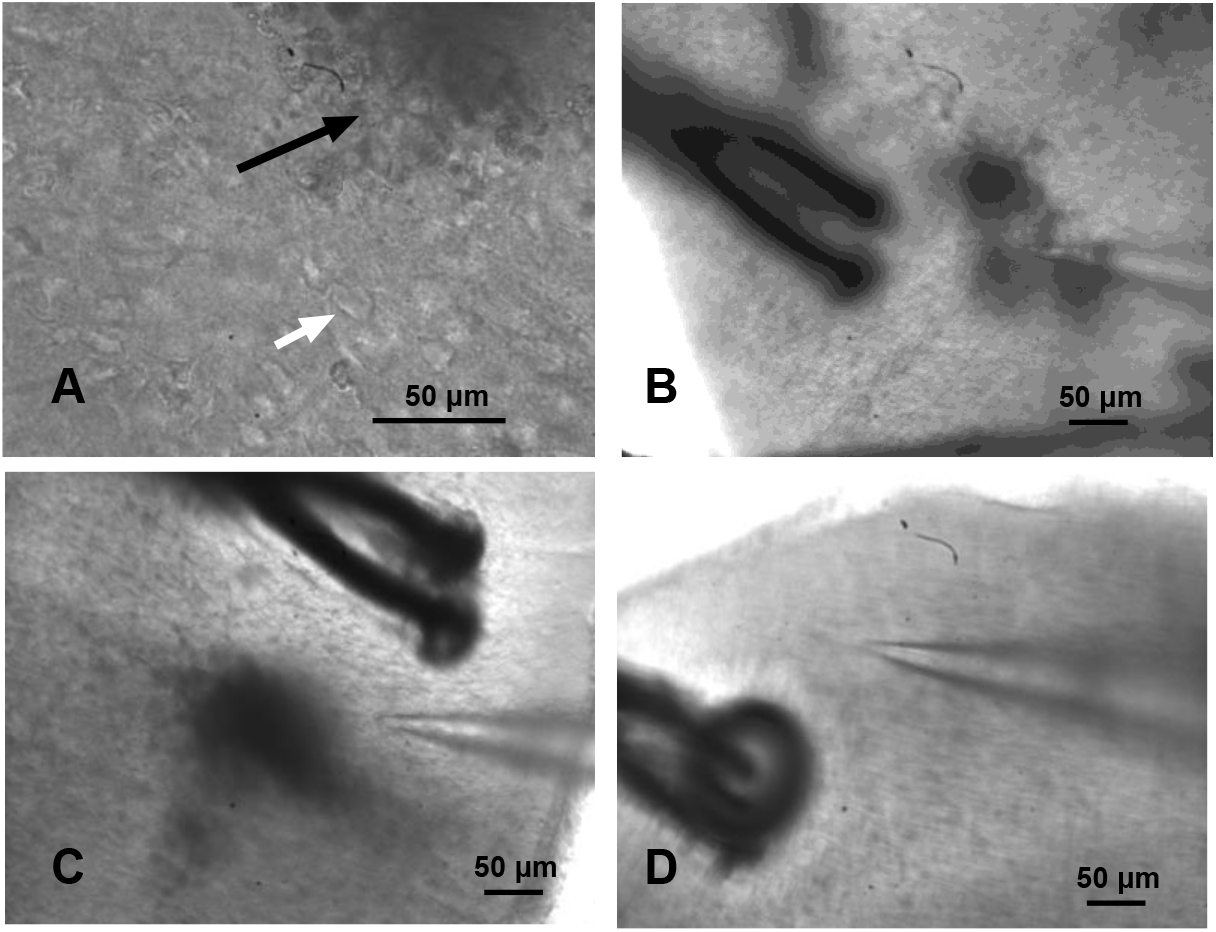
Appearance of injury and recording sites in perfused mouse cortex slices visualized using differential interference contrast at 24 hours after neonatal H-I at P9. A. Opaque regions of focal injury (black arrow) due to light scattering in edematous tissue, surrounded by normally appearing cells (white arrow), 40x objective. A glass pipette used for recording field potentials was placed inside the lesions (B), near the lesion’s border (C), distant from the lesion (B), and on the contralateral hemisphere, 10x objective.

The extracellular field responses (Figure 2A) consisted of a short latency volley burst (VB) followed by the first short latency fEPSP, and often by a longer latency second fEPSP. These responses were reproducible at all recorded sites in animals after H-I and in sham controls. The VBs were insensitive to neurotransmitter receptor blockade and were presumed to be presynaptic. The later components, the first and second field potentials, were completely blocked by 50 µM CNQX and reflected monosynaptic (FP1, latency 4-6 ms) and polysynaptic activity (FP2, latency over 6-10 ms), both mediated by glutamate via AMPA receptors [11]. No other components, such as NMDA, kainate, or GABAergic transmission components, appeared in the field potentials after CNQX and PTX blockade. VBs were entirely eliminated by blocking axonal transmission with 1 µM Tetrodotoxin (TTX).

Input-output fEPSP curves, obtained in response to different levels of stimulation, are shown for FP1 and FP2 in Figures 3B and 3C. Due to the non-linear shape of the curves, with a slight plateau observed in the stimulation range between 0 and 650 µA, the areas under each input-output curve were calculated as a quantitative measure of synaptic strength and neuronal excitability.

**Figure 3.**
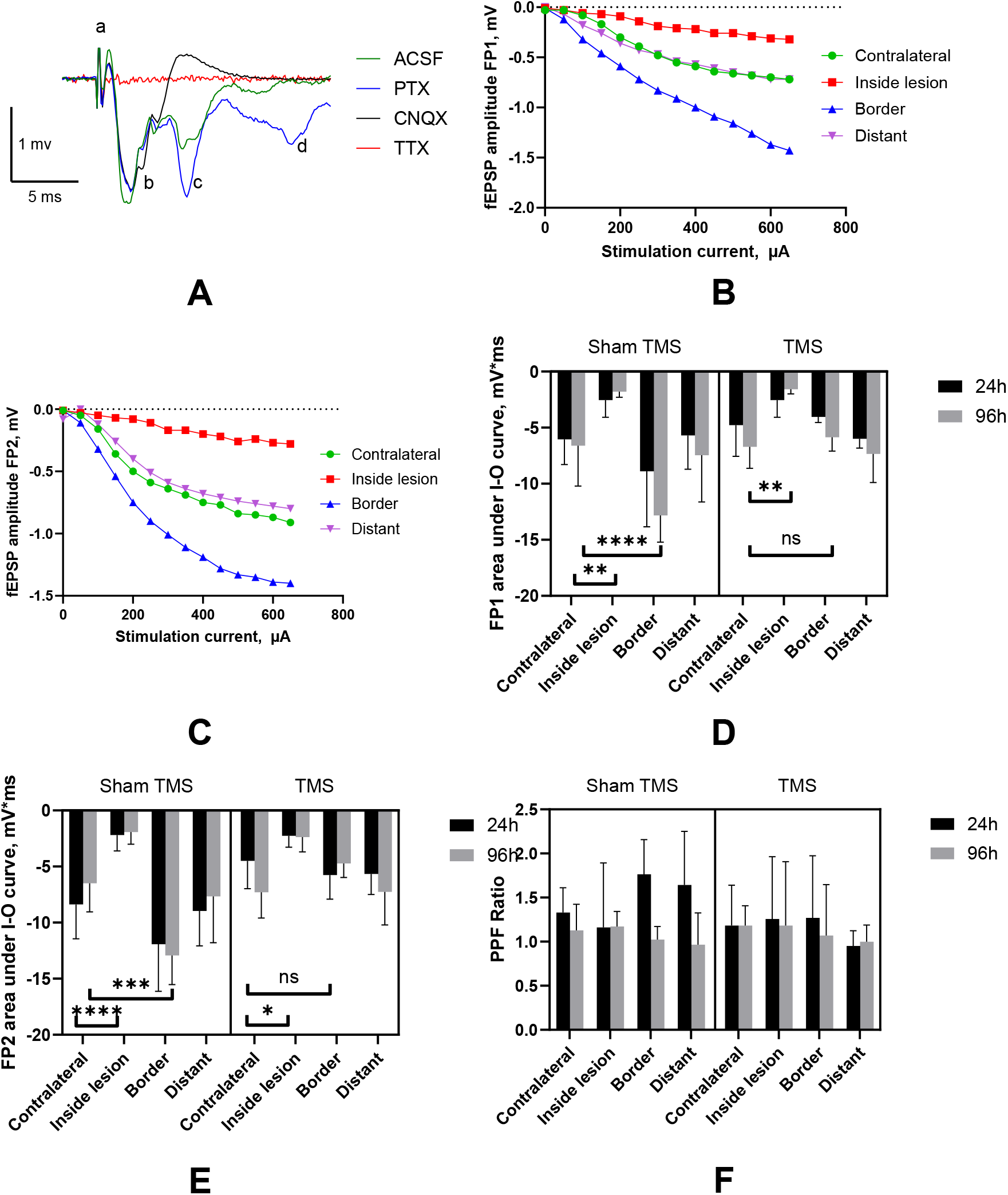
Recording of fEPSP in layer 3 of mouse motor cortex slices after neonatal hypoxia-ischemia. A. Examples of fEPSP and waveforms originating in the P10 mouse cortex. A - Truncated stimulation artifact, b – volley burst, c–First monosynaptic fEPSP, d - Second polysynaptic fEPSP. The legend below shows inhibitors used to identify the origin of components. B, C Input-output curves for the field potential with the short latency (FP1, panel B) and the longer latency (FP2, panel C) in the mouse cortex at proximal and distant sites relative to the edema lesion 24 hours after H-I. The slope of the I-O curve was larger in the lesion border areas for FP1 and FP2. D, E – 3-way ANOVA with post-hoc comparisons revealed a significant increase in the areas under the I-O curves in lesion border areas compared to the contralateral side, and this increase became insignificant with TMS treatment. F-No difference in paired pulse facilitation was observed between the treatment groups and recording locations. **-p<0.01, ****-p<0.0001

### Early and persistent increase of glutamatergic synaptic transmission on the border of the edematous region after H-I

A 3-way ANOVA revealed that changes in the monosynaptic glutamatergic response FP1, measured as an area under the I-O fEPSP curve, depended on the recording location, F(3,112)=31.8, p<0.0001, and the timing of the recording, F(1,112)=13.56, p=0.0004, both of which were significant factors. Compared to normal tissue, lower amplitudes of the fEPSPs and reduced excitability on I-O curves were observed within the edematous regions (post-hoc comparison with contralateral hemisphere p<0.0001). Conversely, potentiation of the monosynaptic glutamatergic response FP1 was detected at the border area (Figure 2B, D, p=0.009) as early as 24 hours after H-I. Additionally, there was a significant interaction between location and recording time in the area under the FP1 I-O fEPSP curve, F(3,112)=10.54, p<0.0001.

A similar increase was observed in polysynaptic (Figure 3 C, E) glutamatergic monosynaptic FP2 responses at the border of edematous regions (post-hoc comparison with the contralateral hemisphere p<0.0001), with significant effects of recording location, F(3,112)=40.93, p<0.0001, and recording time, F(1,112)=30.02, p<0.0001, as shown by ANOVA. This potentiation also lasted for at least 96 hours after H-I. Additionally, there was a significant interaction between location and recording time in the area under the FP2 I-O fEPSP curve, F(3,112)=12.582, p<0.0001.

No differences were found between monosynaptic FP1 and polysynaptic FP2 in the remote recording sites away from the edematous lesions, compared to the contralateral hemisphere. Pair pulse facilitation was slightly higher in sites on the border and away from the edematous lesion (Figure 3F). However, ANOVA did not show a significant main effect of factors, location, or timing of recording after H-I.

### Low-frequency TMS reduces synaptic potentiation following H-I

Following low-frequency transcranial magnetic stimulation (TMS) treatment 30 minutes after H-I, a reduction in potentiated glutamatergic monosynaptic FP1 and polysynaptic responses FP2 at the border of the edematous lesions was observed (p=0.015 at 24 hours and p<0.001 at 96 hours), restoring them to control levels on the contralateral side (Figure 2Bd, E). ANOVA revealed a significant effect of TMS treatment, F(1,112)=7.36, p=0.00077, and an interaction between TMS and recording location factors, F(1,112)=3.06, p=0.033 for FP1. The TMS treatment effect was not significant in the ANOVA for FP2 and PPF. TMS did not affect a decrease in fEPSP responses at either the inside or distant sites of the edematous lesions.

### Low-frequency TMS did not affect long-term cortical neuronal excitability in sham normoxic mice

To determine if 30-minute low-frequency TMS exposure causes long-lasting changes in neuronal excitability in the normal mouse cortex, we compared the area under the I-O fEPSP curve in the motor cortex of sham normoxic mice and the contralateral hemisphere to the ligation in mice after H-I, 24 hours post-TMS or sham TMS exposure, with n = 8 mice per group. Two-way ANOVA did not show a significant effect of either TMS (F(1.28)=1.89, ns) or H-I (F(1,28)=0.54, ns), nor their interaction, on the I-O curve area of the fEPSPs.

### Evolution of edema volume following H-I in the neonatal brain

Cytotoxic edema was easily visible as hyperintense areas on T2-weighted MRI as early as 2 to 4 hours after H-I (Figure 4A). The resolution of cytotoxic edema over time varied between animals but followed a similar pattern in TMS treatment groups, with the most significant edema observed at 24 hours post-H-I and a gradual reduction in edema volume by the end of the second week, along with the formation of cystic lesions and/or tissue loss in the ipsilateral hemisphere. Edematous regions on T2-weighted MRI were further divided by injury severity into hypoxic-ischemic core and penumbra based on regional differences in T2 times in histograms, as described in Methods (Figure 4 B, C). Tissue loss not explained by the ischemic core and penumbra was calculated by subtracting the volumes of the contralateral and injured ipsilateral hemispheres (Figure 4D). This was most noticeable after a week of H-I. The temporal progression of total injury volumes, assessed on T2-weighted MRI, is shown in Figure 4E for mouse pups with and without TMS treatment. In both groups, the peak of total injury occurred at 24 hours post-H-I, with a gradual decrease in the core and penumbra regions and a slight increase in total injury volume by the second week after H-I. The passage of time significantly influenced changes in ischemic core volumes (F(2.207,19.86)=9.329, p=0.0011) and penumbra volumes (F(2.131,19.18)=28.19, p<0.0001) in the repeated measures two-way ANOVA.

**Figure 4.**
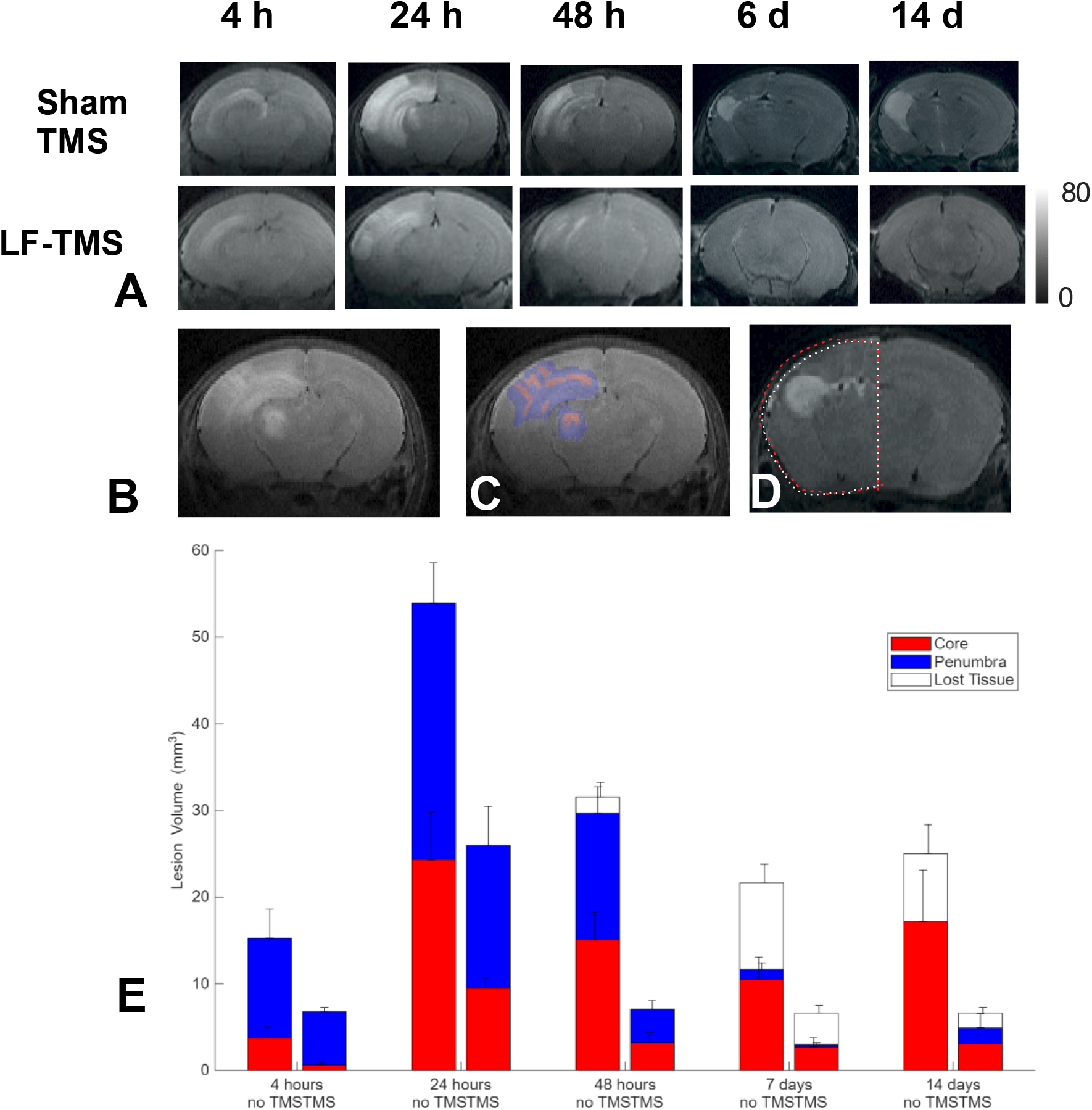
Evolution of hypoxic-ischemic core and penumbra of the edematous lesion after neonatal hypoxia-ischemia and low-frequency TMS treatment. A. Appearance of edematous lesions on T2 time parametric maps at 4, 24, 46 hours, 7, and 14 days after H-I. Edematous regions (B) were stratified into hypoxic-ischemic core and penumbra (C, red and blue). Lost tissue was calculated by subtracting brain areas in the ipsilateral and symmetrical contralateral hemispheres (D). E. Changes in ischemic core, penumbra, and lost tissue classes over time in sham TMS and LF-TMS groups. Data are presented as means ± standard deviation.

### Low-frequency TMS reduces the volume of the hypoxic-ischemic core and penumbra after H-I

LF-TMS treatment significantly altered the volumes and proportions of the core (F(1,9)=9.302, P=0.0138) and penumbra (F(1,9)=5.307, P=0.0467) during the H-I edematous injury resolution process, as analyzed with a repeated measures two-way ANOVA. Post hoc comparison of mean volumes at each studied time point showed a significant decrease in the core (p=0.043) and a non-significant reduction in the penumbra (p=0.367) in the LF-TMS group at 48 hours after H-I. The interaction of Time and TMS treatment factors was also significant for the core (F(2.207,19.86)=1.911, P=0.1714) and the penumbra (F(2.131,19.18)=3.400, P=0.0519). No significant differences between the TMS treatment groups were observed in the lost tissue class.

### Expression of the GluR2 subunit in the cortex penumbra after H-I or TMS

Given the previously reported role of the AMPA receptor subunit GluR2 in regulating calcium permeability of AMPA receptors and exacerbating injury in global and focal hypoxia in adult [12] and neonatal [13] animal models, we examined Gria2 gene expression, which encodes the GluR2 subunit of AMPA receptors, in the somatosensory cortex bordering the edematous lesion core and penumbra, guided by T2-weighted MRI in P11-P12 mice. Two-way ANOVA reveals a significant effect of H-I (F(2,49)=4.016, p=0.024), and there was a decrease in Gria2 expression in the ipsilateral cortex after H-I compared to sham H-I on post-hoc multiple comparisons (p=0.0238), Figure 5. There was no difference between those groups after TMS treatment.

**Figure 5.**
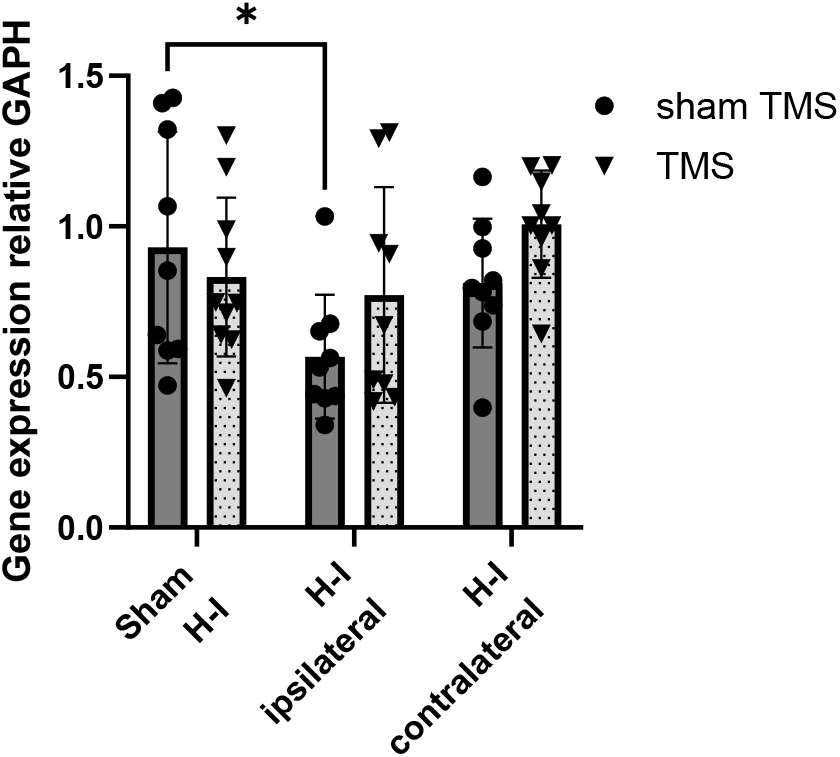
Gene expression on PCR in the somatosensory cortex penumbra at P11-P12 after H-I at P9. Two-way ANOVA reveals a significant effect of H-I, and there is a decrease in Gria 2 expression in the ipsilateral cortex after H-I compared to sham H-I.

## DISCUSSION

Our study demonstrates that low-frequency transcranial magnetic stimulation (LF-TMS) provides both acute and delayed neuroprotection in a neonatal hypoxic-ischemic (H-I) mouse model. A single session of LF-TMS, administered shortly after H-I, significantly reduces synaptic potentiation and mitigates the volume of injured brain tissue. This suggests that LF-TMS could serve as a promising adjunct therapy to augment and address the limitations of therapeutic hypothermia, which is currently the only FDA-approved treatment for neonatal HIE but does not offer complete protection [3].

### Hypoxic Long-Term Potentiation

Our electrophysiology results indicate that H-I leads to a pathological increase in glutamatergic transmission, which is evident as a potentiation of both monosynaptic and polysynaptic fEPSP responses in brain regions bordering the edematous lesion. We, along with other researchers, have observed that a sustained pathological increase in glutamatergic transmission is a key mechanism of perinatal brain injury [6, 14]. This phenomenon was observed in vivo and ex vivo in neonatal intermittent hypoxia model [6], ex vivo hypoxia models [5], and termed this effect ‘post-hypoxic long-term potentiation (hLTP)’. This potentiation was driven by AMPA and NMDA receptor activation and increased calcium influx, with no change in paired-pulse facilitation, engaging similar mechanisms to classic LTP, in addition to more AMPA receptors on the post-synaptic surface. This makes the synapse more sensitive and increases the likelihood of firing an action potential.

Our study adds a crucial finding: this hypoxic potentiation was specifically observed in the hypoxic penumbra, the brain areas bordering the ischemic core. We believe this is significant because this may increase neuronal activity, glutamate release, and excitotoxicity, leading to further brain injury in this vulnerable region. Hyperexcitability of neurons is likely a key contributor to seizure activity and neuronal damage in neonates after perinatal H-I [15, 16].

### Proposed Therapeutic Mechanism of LF-TMS

Drawing on the similarities between the cellular mechanisms of classical LTP and hLTP [5], we suggest that low-frequency transcranial magnetic stimulation (LF-TMS) can serve as a targeted intervention to reverse the abnormal potentiation of synapses. This works through mechanisms similar to long-term depression (LTD), a form of synaptic plasticity that weakens neuronal connections. Low-frequency electric stimulation has been successfully used to restore elevated fEPSP to control levels in the brain after cardiac arrest and restore LTP in neonatal intermittent hypoxia [6]. LF-TMS has been extensively used to reduce cortical excitability in stroke [17]. Our study found that a single session of LF-TMS successfully returned the heightened glutamatergic responses, a key indicator of hLTP, to control levels. While not affecting receptor production, LF-TMS is believed to trigger a cellular cascade that leads to the post-translational regulation of AMPA receptor redistribution. This mechanism effectively removes existing AMPA receptors from the postsynaptic membrane, reducing the synapse’s sensitivity to glutamate and decreasing excitotoxicity in the penumbra—the brain region bordering the ischemic core.

This finding supports the concept of hypoxic synaptic potentiation as a key mechanism contributing to excitotoxicity after perinatal brain injury. The fact that this potentiation can last for at least 96 hours after H-I emphasizes its role as a prolonged pathological process that needs intervention. Importantly, LF-TMS treatment effectively brought these enhanced glutamatergic responses back to control levels, indicating that TMS can adjust abnormal excitability in the injured neonatal brain. The observation that LF-TMS did not change excitability in sham-treated, normoxic mice confirms its safety profile and supports its use as a focused intervention for pathological conditions without impacting normal brain function. This is a significant advantage over systemic drug administration, which can disrupt essential neurodevelopmental signaling pathways.

The in-vivo MRI results support the neuroprotective effects observed with electrophysiology. Serial MRI scans demonstrated that LF-TMS significantly decreased the volumes of both the ischemic core and penumbra at 48 hours post-H-I. This reduction in injured tissue volume is a key outcome, as it directly relates to the long-term preservation of brain function. Although the study did not find a significant effect on the long-term “lost tissue” class, the decrease in acute cytotoxic edema is an essential step toward limiting permanent brain damage. The early and substantial effect on cytotoxic edema indicates that LF-TMS may be most effective when given within a narrow therapeutic window shortly after the H-I event.

The study also examined the role of the GluR2 subunit, which is known to regulate calcium permeability in AMPA receptors and has been linked to worsening H-I injury in animal models [12]. Although previous research highlighted the contribution of the GluR2 subunit to worsening H-I injury, our findings show that the neuroprotective effects of LF-TMS are not solely due to changes in GluR2 expression. The results reveal that H-I causes a reduction in the Gria2 gene expression—which encodes the GluR2 subunit—in the ipsilateral cortex compared to sham controls. This aligns with existing literature demonstrating that decreased GluR2 expression leads to the formation of calcium-permeable AMPA receptors, a key mechanism involved in neuronal death in models of transient forebrain ischemia and perinatal brain injury [12, 13]. Our neonatal H-I data support this well-established mechanism. However, the effect of TMS on Gria2 expression was not definitive. While there was no significant difference in Gria2 expression between the H-I group and the H-I group treated with TMS, there was also no significant difference between the TMS group and the sham control group. These findings, while encouraging, remain inconclusive regarding the role of Gria2, and additional experiments at different time points are needed to clarify the exact mechanism of TMS action.

There could be other possible mechanisms for the neuroprotective action of TMS in acute hypoxic events in the brain. A recent study in an adult rat focal stroke model observed that the neuroprotective effect of LF-TMS was due to upregulation of the antiapoptotic factor Bcl-2 and downregulation of the proapoptotic caspase-3 cleavage, as well as decreased expression of Bax and matrix metallopeptidase-9 [18].

In summary, this study offers strong evidence that LF-TMS is a promising, non-invasive neuromodulation method for treating neonatal HIE. By effectively reducing harmful synaptic potentiation and decreasing the size of acute injuries, it could serve as a valuable supplement to therapeutic hypothermia, opening new opportunities for improving neurodevelopmental outcomes in affected newborns. Future research is needed to understand the exact molecular and cellular mechanisms of LF-TMS and to assess its long-term effects on functional recovery and cognitive development.

## Acknowledgments

This study was funded by NIH grants 1R01NS119251-01A1

## REFERENCES

1. Gunn, A.J., Cerebral hypothermia for prevention of brain injury following perinatal asphyxia. Current Opinion in Pediatrics, 2000. 12(2): p. 111–5.

2. Davidson, J.O., et al., Therapeutic Hypothermia for Neonatal Hypoxic-Ischemic Encephalopathy -Where to from Here? Front Neurol, 2015. 6: p. 198.

3. Cilio, M.R. and D.M. Ferriero, Synergistic neuroprotective therapies with hypothermia. Semin Fetal Neonatal Med, 2010. 15(5): p. 293–8.

4. Ovcjak, A., et al., Hypothermia combined with neuroprotective adjuvants shortens the duration of hospitalization in infants with hypoxic ischemic encephalopathy: Meta-analysis. Front Pharmacol, 2022. 13: p. 1037131.

5. Heit, B.S., et al., Interference with glutamate antiporter system x(c) (-) enables post-hypoxic longterm potentiation in hippocampus. Exp Physiol, 2024. 109(9): p. 1572–1592.

6. Goussakov, I., et al., Occlusion of activity dependent synaptic plasticity by late hypoxic long term potentiation after neonatal intermittent hypoxia. Exp Neurol, 2021. 337: p. 113575.

7. Novak, I., et al., Early, Accurate Diagnosis and Early Intervention in Cerebral Palsy: Advances in Diagnosis and Treatment. JAMA Pediatr, 2017. 171(9): p. 897–907.

8. Zhu, C., X. Wang, and K. Blomgren, Cerebral Hypoxia—Ischemia in Neonatal Rats or Mice: A Model of Perinatal Brain Injury, in Animal Models of Acute Neurological Injuries, J. Chen, et al., Editors. 2009, Humana Press: Totowa, NJ. p. 221–230.

9. Boonzaier, J., et al., Design and Evaluation of a Rodent-Specific Transcranial Magnetic Stimulation Coil: An In Silico and In Vivo Validation Study. Neuromodulation, 2020. 23(3): p. 324–334.

10. Drobyshevsky, A., et al., Functional correlates of central white matter maturation in perinatal period in rabbits. Exp Neurol, 2014. 261C: p. 76–86.

11. Wallace, J., et al., Characterization of electrically evoked field potentials in the medial prefrontal cortex and orbitofrontal cortex of the rat: modulation by monoamines. Eur Neuropsychopharmacol, 2014. 24(2): p. 321–32.

12. Liu, S., et al., Expression of Ca(2+)-permeable AMPA receptor channels primes cell death in transient forebrain ischemia. Neuron, 2004. 43(1): p. 43–55.

13. Jensen, F.E., The role of glutamate receptor maturation in perinatal seizures and brain injury. International Journal of Developmental Neuroscience, 2002. 20(3): p. 339–347.

14. Jensen, F.E., et al., Acute and chronic increases in excitability in rat hippocampal slices after perinatal hypoxia In vivo. J Neurophysiol, 1998. 79(1): p. 73–81.

15. Burnsed, J., et al., Increased glutamatergic synaptic transmission during development in layer II/III mouse motor cortex pyramidal neurons. Cereb Cortex, 2023. 33(8): p. 4645–4653.

16. Burnsed, J., et al., Neuronal Circuit Activity during Neonatal Hypoxic–Ischemic Seizures in Mice. Annals of Neurology, 2019. 0(0).

17. Wang, C., et al., Effects of Low-Frequency (0.5 Hz) and High-Frequency (10 Hz) Repetitive Transcranial Magnetic Stimulation on Neurological Function, Motor Function, and Excitability of Cortex in Ischemic Stroke Patients. Neurologist, 2023. 28(1): p. 11–18.

18. Buetefisch, C.M., et al., Neuroprotection of Low-Frequency Repetitive Transcranial Magnetic Stimulation after Ischemic Stroke in Rats. Ann Neurol, 2023. 93(2): p. 336–347.

